# Astrocyte and mitochondrial footprints in brain-derived extracellular vesicles predict tau pathology

**DOI:** 10.1101/2025.01.09.632249

**Authors:** Jeanne Espourteille, Aatmika Barve, Valentin Zufferey, Elodie Leroux, Romain Perbet, Séverine Bégard, Raphaëlle Caillierez, Claude-Alain Maurage, Nicolas Toni, Luc Buée, Morvane Colin, Kevin Richetin

**Author notes:** These authors contributed equally to this work (Jeanne Espourteille, Aatmika Barve). Correspondence to: Kevin Richetin Full address: Department of Psychiatry, Center for Psychiatric Neurosciences, Lausanne University Hospital (CHUV) and University of Lausanne, 1008 Prilly-Lausanne, Switzerland.

## Abstract

Tauopathies are neurodegenerative disorders characterized by abnormal tau aggregation, with primary 3R (e.g., Pick’s disease, PiD) and 4R (e.g., progressive supranuclear palsy, PSP) variants posing a significant diagnostic challenge. Here, we examined brain-derived extracellular vesicles (BD-EVs) isolated from the prefrontal cortex of PiD (3R), PSP (4R), and non-demented controls (CTRL) to determine if these vesicles reflect disease-specific proteomic signatures.

We found that while tau pathology does not substantially alter BD-EV concentration or the enrichment of core vesicular markers, it does influence their size distribution and protein cargo. BD-EV samples from PiD patients exhibited a greater abundance of small vesicles and distinct protein profiles when compared to PSP and CTRL. Weighted Gene Co-expression Network Analysis (WGCNA) identified four key protein modules to account for variance between patient groups Endoplasmic Reticulum, Mitochondria, Microtubules, and Trivalent Inorganic Cation Transport. In PiD, astrocyte-derived mitochondrial proteins were significantly elevated, whereas neuronal microtubule-related proteins were diminished relative to both PSP and CTRL. Notably, changes in the mitochondrion and microtubule modules enhanced the detection of PiD pathology. Cellular origin annotation revealed a marked shift in BD-EV composition: PiD samples exhibited an increased astrocytic signature, while both PiD and PSP showed a reduction in neuronal proteins compared to CTRL. Crucially, the enrichment of astrocytic mitochondrial and endoplasmic reticulum proteins, alongside reduced neuronal proteins, correlated strongly with the severity of tau pathology (AT8-stained aggregates) in patient brains.

These findings demonstrate that BD-EVs capture tau isoform-specific cellular and molecular alterations, offering a window into disease mechanisms at the neuron-glia interface. By linking distinct protein signatures and their cellular origins to tau pathology severity, our results highlight the potential of BD-EV profiling as a biomarker strategy for distinguishing between and monitoring the progression of 3R and 4R tauopathies.

## Introduction

Tau protein is pleiotropic, performing multiple essential roles in the brain. It stabilizes microtubules, ensuring efficient nutrient transport and cellular communication along axons. Additionally, tau plays a crucial role in synaptic plasticity by modulating neuronal signaling pathways and regulating interactions between various proteins, which are vital for maintaining neural network stability and adaptability ^1–4^.

However, when tau becomes dysfunctional, it gives rise to a class of neurodegenerative diseases known as tauopathies^5^. The heterogeneity of tauopathies is striking; they manifest in a spectrum of clinical presentations and neuropathological profiles. Each tauopathy has distinct clinical and pathological features but shares the commonality of tau dysfunction. The mechanisms leading to tau aggregation are complex and not yet fully understood. They involve abnormal phosphorylation of tau, which reduces its affinity for microtubules and increases its propensity to form intracellular aggregates ^6,7^. The specific pattern of tau deposition, the isoforms of tau which aggregate, and the regional brain involvement vary across different tauopathies, which can influence the clinical presentation and progression of the disease ^5,8–10^. For instance, Pick’s disease (PiD) is typically characterized by the presence of "Pick bodies", inclusions of 3R tau in granular neurons of the dentate gyrus of the hippocampus and layers two and six of the prefrontal cortex, whereas progressive supranuclear palsy (PSP) and corticobasal degeneration (CBD) are characterized by neuronal and glial ("tufted" astrocytes and astrocytic plaques) lesions made of 4R tau in the brainstem, cerebellar, subcortical and cortical brain regions ^11,12^. This diversity in tauopathies suggests that while tau is a common pathological thread, disease-specific factors determine the more subtle phenotypes of this family of diseases ^13^.

Clinicians today still struggle to differentiate and diagnose tauopathies early in the disease progression. This is particularly true of primary tauopathies, for which the main protein driver is tau. Circulating biomarkers like phosphorylated tau (P-tau) in blood and cerebrospinal fluid (CSF) are increasingly accurate at reflecting tau pathology in the brain at earlier time points in disease progression, but they are often only validated for pathologies like AD which have amyloid beta as a co-driver of pathology ^14^. Some tau positron emission tomography (PET) tracers can differentiate 4R tau inclusions and thus can identify diseases like PSP and CBD, however, these tracers are not yet validated for clinical use ^15,16^.

Interestingly, proteomic analysis of tau inclusions in human 3R and 4R pathology is capable of distinguishing specific tauopathies based on post-translational modifications present in the insoluble tau aggregates obtained from brain tissue. While insoluble tau exhibited distinct phenotypes specific to each pathology, soluble tau extracted from the same aggregates was unable to differentiate between these pathologies ^17^. Various forms of soluble hyperphosphorylated tau are currently measured in blood and CSF for diagnostic purposes, but to date they do not allow the diagnosis of primary 3R and 4R tauopathies ^18^.

Extracellular vesicles (EVs) are now a promising diagnostic tool for many brain diseases. Brain-derived EVs (BD-EVs), which are shed by cells into the interstitial spaces of the brain, encapsulate a diverse cargo that reflects the physiological (or pathological) state of their parent cells ^19^. This encapsulated cargo includes proteins, lipids, RNA, and other molecules, providing a comprehensive snapshot of the cellular environment at any given time ^20^. As such, BD-EVs provide a unique window into brain health, revealing information that is otherwise difficult to obtain. Thanks to their proteomic cargo, BD-EVs could be the key to a better understanding of pathophysiology, and if their content is also available in accessible biofluids such as plasma and saliva, they could be a diagnostic tool for various tauopathies. This potential has not gone unnoticed, as numerous researchers are exploring using BD-EVs as biomarkers for a wide range of neurodegenerative disorders ^21^. Recently, Chatterjee and colleagues have demonstrated that the 3R/4R ratio of plasmatic EV isolated by L1CAM could serve as a suitable diagnostic tool to discriminate different forms of frontotemporal dementia, ALS and Parkinson’s disease ^22^.

Nevertheless, little is known about the protein signature of BD-EVs as a function of the different types of tau inclusions and isoforms. In this study, we explored the potential of BD-EVs to enhance the understanding and diagnosis of 3R and 4R tauopathies. We analyzed the proteomic content of BD-EVs isolated from the prefrontal cortical biofluids of patients with different tauopathies, comparing their cellular and molecular signatures. Using a combination of advanced approaches, we aimed to identify specific biomarkers that could reflect the severity and underlying mechanisms of tauopathies. Combined with current diagnostic protocols, these markers could pave the way for more accurate diagnosis and dementia classification, particularly for non-AD tau pathology.

## Materials and methods

### Patient cohort

Brain extracts were obtained from Lille Neurobank, fulfilling requirements concerning biological resources and declared to the competent authority. Samples were managed by the CRB/CIC1403 Biobank (BB-0033-00030 for Lille Neurobank). The demographic data are listed in Leroux et al. ^23^.

### Interstitial fluid isolation from the patient prefrontal cortex tissue

Brain derived fluids (BD-fluids) were isolated as previously described ^24^ Briefly, roughly 1.5g of fresh-frozen patient prefrontal cortex were gently cut into smaller pieces. These were then incubated in a Papain/Hibernate-E solution to gently digest the tissue and release interstitial fluid, theoretically avoiding cellular lysis. After tissue digestion, differential centrifugations at 300g for 10 min, 2,000g for 10 min, and 10,000*g* for 30 min were performed at 4°C to remove cells, membranes, and debris, respectively, and the supernatant of the 10,000*g* centrifugation was then stored at -80°C until EV isolation was performed. Normalization according to the weight of the brain extracts was systematically done to avoid bias in the results.

### Patient brain-derived extracellular vesicle isolation from interstitial fluid

The procedures to isolate the brain derived EVs (BD-EVs) from the human BD-fluid were carried out in accordance with the minimal information for the studies of extracellular vesicles (MISEV) guidelines that were established and updated in 2024 by the International Society for Extracellular Vesicles ^25^. Various controls were applied to validate the enrichment and the content of the BD-EVs, as recommended in these guidelines. However, the procedure described above to recover BD-fluids may still lead to some cell lysis and intracellular contamination. Intraluminal vesicles (ILVs) in the preparations cannot be fully excluded. 500 μL of BD-fluid were loaded on the top of a size exclusion chromatography (SEC) column (10mL column, CL2B Sepharose, pore size 75 nm, Millipore), ^26^. A mean of 7.94 x 10^10^ (±3.36 x 10^9^ vesicles/g of tissue in F1–4 (n = 36 samples) was recovered for each sample. Isolation was carried out in phosphate-buffered saline (PBS) with a flow of 36–48 s/mL. The first 3mL were eliminated, and the following 20 fractions were recovered (500 μL per fraction).

### Measurement of brain derived extracellular vesicle size and concentration

Nanoparticle tracking analysis (NTA) was performed on individual fractions diluted in PBS with a NanoSight NS300 (Malvern Panalytical). To generate statistical data, five videos of 90 s were recorded and analyzed using NTA software (camera level: 15; detection threshold: 4).

### Label-free liquid chromatography tandem mass spectrometry

#### Protein digestion

F1-4 fractions were digested according to a modified version of the iST method (named miST method) ^27^. Briefly, 50mL solution in PBS were supplemented with 50mL miST lysis buffer (1% Sodium deoxycholate, 100 mM Tris pH 8.6, 10 mM DTT) and heated at 95°C for 5 min. Samples were then diluted 1:1 (v:v) with water and reduced disulfides were alkylated by adding 1 /4 vol of 160 mM chloroacetamide (final 32 mM) and incubating at 25°C for 45 min in the dark. Samples were adjusted to 3 mM EDTA and digested with 0.5mg Trypsin/LysC mix (Promega #V5073) for 1h at 37°C, followed by a second 1h digestion with a second and identical aliquot of proteases. To remove sodium deoxycholate and desalt peptides, two sample volumes of isopropanol containing 1% TFA were added to the digests, and the samples were desalted on a strong cation exchange (SCX) plate (Oasis MCX; Waters Corp., Milford, MA) by centrifugation. After washing with isopropanol/1% TFA, peptides were eluted in 250 mL of 80% MeCN, 19% water, 1% (v/v) ammonia.

#### Liquid chromatography-tandem mass spectrometry

Tryptic peptide mixtures were injected on an Ultimate RSLC 3000 nanoHPLC system interfaced via a nanospray Flex source to a high resolution Orbitrap Exploris 480 mass spectrometer (Thermo Fisher, Bremen, Germany). Peptides were loaded onto a trapping microcolumn Acclaim PepMap100 C18 (20mm x 100μm ID, 5μm, Dionex) before separation on a C18 custom packed column (75μm ID × 45cm, 1.8μm particles, Reprosil Pur, Dr. Maisch), using a gradient from 4 to 90% acetonitrile in 0.1% formic acid for peptide separation (total time: 140 min). Full MS survey scans were performed at 120,000 resolution. A data-dependent acquisition method controlled by Xcalibur software (Thermo Fisher Scientific) was used that optimized the number of precursors selected ("top speed") of charge 2+ to 5+ while maintaining a fixed scan cycle of 2s. Peptides were fragmented by higher energy collision dissociation (HCD) with a normalized energy of 30% at 15’000 resolution. The window for precursor isolation was of 1.6 m/z units around the precursor and selected fragments were excluded for 60s from further analysis.

#### MS and MS data analysis

Data files were analyzed with MaxQuant 1.6.14.0 incorporating the Andromeda search engine ^28,29^. Cysteine carbamidomethylation was selected as fixed modification while methionine oxidation and protein N-terminal acetylation were specified as variable modifications. The sequence databases used for searching were the human (Homo sapien) reference proteome based on the UniProt database and a "contaminant" database containing the most usual environmental contaminants and enzymes used for digestion (keratins, trypsin, etc.). Mass tolerance was 4.5 ppm on precursors (after recalibration) and 20 ppm on MS/MS fragments. Both peptide and protein identifications were filtered at 1% FDR relative to hits against a decoy database built by reversing protein sequences.

### Categorical annotation of proteins

Proteins were annotated according to several databases. These databases included the Gene Ontology (GO) Human database - further split into the categories of cellular component (CC), biological process (BP), and molecular functions (MF) - as well as the MISEV guidelines for EV-associated and potential contaminant proteins, a database generated by McKenzie and colleagues in their publication for brain cell type specificity, and the Human Protein Atlas databases for cell type specificity and predicted subcellular localization ^25,30–33^. We also compared excitatory and inhibitory neuron-specific proteins within the neuronal category according to the Human Protein Atlas ^33^. After annotation, according to the McKenzie et al. database, glial abundance was determined by summing the abundance of astrocytic, oligodendrocytic, and microglial material ^32^.

### Weighted gene co-expression network analysis

Protein co-expression network analysis was performed with the R package WGCNA on the preprocessed proteomic data of all identified proteins with their respective riBAQ scores for each disease group ^34^. At first, a correlation matrix for all pair-wise correlations of proteins across all samples was generated and then transformed into a matrix of connection strengths, i.e., a weighted adjacency matrix with soft threshold power β = 16. Then, the topological overlap (TO) was calculated using the connection strengths. Proteins were further hierarchically clustered using 1-TO measure as the distance measure to generate a cluster dendrogram, and modules of proteins with similar co-expression relationships were identified by using a dynamic tree-cutting algorithm with parameters set to minimal module size = 30, deepSplit = 2, and merge cut height = 0.15. Within each module, the module eigenprotein was defined as the first principal component, which serves as a weighted summary of protein expression within the module and accounts for the greatest variance among all module proteins. Further, module membership (kME) was assigned by calculating Pearson’s correlation between each protein and each module eigenprotein and the corresponding p-values. Proteins were reassigned to the module for which they had the highest module membership with a reassignment threshold of *p* < 0.05. Hub proteins were found from each module using the signedkME function, which explains the protein membership with its module and its strong association within the module.

### Gene ontology

A detailed genetic annotation of each protein within WGCNA modules was performed. These differentially expressed proteins and co-expressed proteins were characterized based on their gene ontologies, using GO Elite (version 1.2.5) python package ^35^. The entire set of proteins identified and included in the network analysis served as the background dataset. The presence of significantly overrepresented ontologies within a module was gauged using a Z-score, while the significance of Z-scores was evaluated using a one-tailed Fisher’s exact test, with adjustments for multiple comparisons via the Benjamini-Hochberg FDR method. Threshold analysis included a Z-score cut off 2, a p-value threshold of 0.05, and a minimum requirement of five genes per ontology before ontology pruning was performed. The best gene ontology term which explains the molecular and cellular function of each module was used to name it.

### Brain cell type enrichment

GeneListFET in R was used to evaluate the cell type specificity of WGCNA module proteins ^36,37^. The cell type specific gene list from RNA-seq in human brain was used ^32^. The total list of identified protein groups was used in background and the cell type specific gene list was filtered for presence in the total proteins list prior to cross-referencing. One tailed Fisher’s exact test was performed and correction for multiple comparisons by the FDR (Benjamini Hochberg) method was applied to evaluate the significance of cell type enrichment of each module We have used the -log10(p value) to represent the significance of cell type association.

### Protein-Protein interaction network

Proteins in WGCNA module are associated to some specific biological entity and functions, and these proteins must also be interacting between modules as well. To understand the interactions between the proteins from different modules, we used SIGNOR 3.0^38^ database as our knowledge bank to get the interactions. We utilized Cytoscape 3.10.3 for the network visualization and transformation. Obtained networks were merged into one that was subsequently curated to keep only potential biomarker protein along with their direct or 1-step neighborhood interaction, and additional protein complexes and phenotypes proposed by SIGNOR. We used different styling tools to represents different characteristics of proteins and their interactions. The interaction network is a reduced version of the whole interactome to highlight the important module proteins and interactions between them.

### Machine learning

Machine learning was used to support the traditional statistical results. The dataset was split into 60% training set and 40% testing set. The split was stratified and performed 42 times with different random seeds. Then, Borderline SMOTE (Synthetic Minority Over-sampling Technique) was performed on the training set to balance the classes before training the model. OneVsRestClassifier model performed upon XGBClassifier model was used with parameters set to multi:softprob for objective and mlogloss for evaluation metric ^39,40^. The model was trained with the training set for the classification of data into three classes for each individual feature. The model was evaluated on the testing set and its performance was assessed using F1-score as the metrics for evaluation. The F1-score for each individual feature for three classes was represented in a heatmap.

### Histology

The prefrontal region of these patients, adjacent to the one used to isolate BD-EVs in this study, was previously quantified for AT8 histology (% of lesions) as part of the publication by Leroux and colleagues ^41^. The values from this quantification were reused to perform correlations with the different protein quantities and module scores presented in Figure 4.

### Statistical analyses

All statistical analyses were performed using GraphPad Prism V9.4. Statistical significance was evaluated using ordinary one-way ANOVA with Tukey’s multiple comparisons post hoc test for particle concentration, MAPT abundance, endoplasmic reticulum protein eigengene plot, and cation transport protein eigengene plot. Ordinary two-way ANOVA was used to evaluate brain specific vs. nonspecific protein abundance. Ordinary two-way ANOVA with Dunnett’s multiple comparisons post hoc test was used to evaluate the proportion of brain specific protein which was brain cell type specific. Ordinary two-way ANOVA with Tukey’s multiple comparisons post hoc test was used to evaluate AT8 and AT100 inclusion analysis. Ordinary two-way ANOVA with Sidak’s multiple comparisons post hoc test was used to evaluate the EV associated protein abundance and neuronal vs. glial protein abundance. Kruskal-Wallis with Dunn’s multiple comparisons test was used for non-normal data analyses, including the quantity of LEV vs. SEV, SNCA abundance, APP abundance, mitochondrial protein eigengene plot, and microtubule protein eigengene plot. R values were plotted in heatmaps for correlation matrices comparing various BD-EV features with histopathological findings. Systematically, a two-tailed Pearson’s correlation test was run with 95% confidence interval. Significance was systematically denoted with stars: * *p* < 0.05, ** *p* < 0.01, *** *p* < 0.001, **** *p* < 0.0001.

## Results

### 1 Tau isoform-specific pathology doesn’t impact the distribution profile of BD-EVs

Prefrontal cortex samples were prepared to isolate brain derived (BD)-fluids according to the protocol described by Leroux and colleagues^23^. Then, BD fluid-derived EVs (BD-EV) were isolated from human BD fluid using differential centrifugations and size exclusion chromatography (SEC). BD-EVs were then subjected to nanoparticle tracking analysis (NTA) to evaluate their size and concentration, as well as liquid chromatography tandem mass spectrometry (LC-MS/MS) to evaluate their protein content (**Figure 1A**). Firstly, measurements with NTA indicated that PiD and PSP pathology do not affect the concentration of BD-EVs circulating in the patient’s prefrontal cortex fluids (**Figure 1B-C**). PiD pathology does, however, shift the size distribution of the EV sample towards having a higher ratio of small EVs (<150nm) to large EVs (>150nm) than CTRL (**Figure 1C-D**). We used proteomic analysis to evaluate the quality of BD-EV isolations. We revealed that most proteins detected in patient BD-EVs are shared among control and pathological samples, while a few proteins are detected only in PiD or PSP **(Figure 1E**). Using the minimum information for the study of EVs (MISEV) guidelines, we annotated all proteins according to their known association with EVs as potential EV markers or potential contaminants of EV isolation ^25^. This analysis revealed a significantly higher abundance of EV-associated proteins than components of non-EV associated co-isolated structures **(Figure 1F**) in all three patient groups, indicating that EV samples are highly enriched in vesicular material. Altogether, these results indicate that tau pathology does not influence the quantity or quality of BD-EVs secreted into patients’ brain derived fluid, but may affect other aspects of BD-EVs’ proteomic makeup and size distribution.

**Figure 1.**
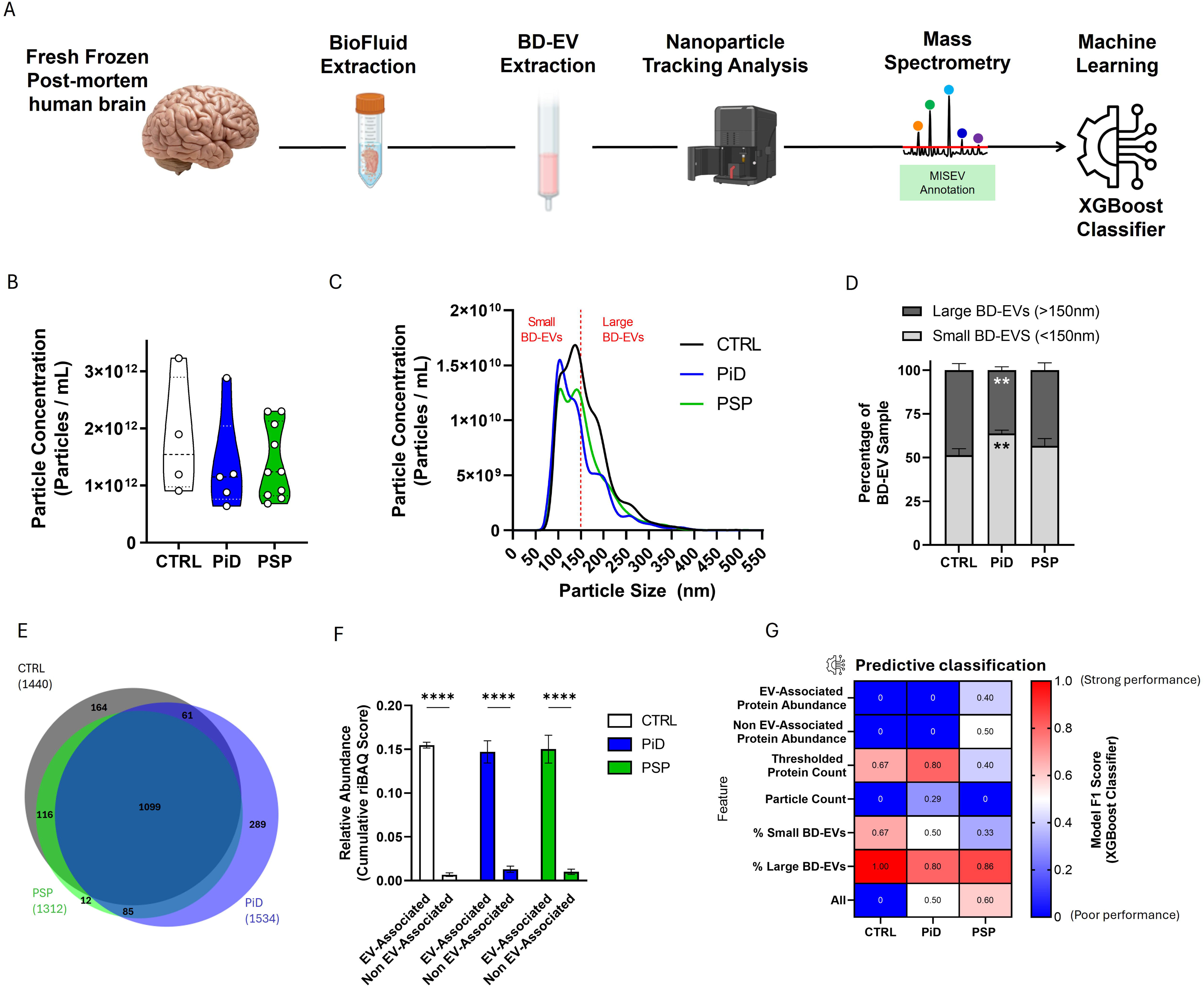
Consequences of tau pathology on concentration, size and quality of BD-EVs. **(A)** Illustration of the procedure to investigate how BD-EVs may reflect brain tau pathology. Highlighted is the MISEV annotation of proteins in BD-EV samples as EV-associated or non-EV-associated. **(B)** Violin plot depicting the concentration of BD-EVs detected by NTA by patient group. **(C)** Area graph depicting the size and number of EVs in each patient group. Particle concentration has been normalized by tissue weight prior to EV extraction. **(D)** Stacked bar graph showing the ratio of small (<150nm) to large (>150nm) EVs by patient group. **(E)** Venn diagram showing the number of proteins present after threshold filtration in the BD-EVs of CTRL, PiD and PSP patients. **(F)** Bar graph quantifying the abundance of protein associated with EVs vs. potentially contaminating proteins of EV isolation according to MISEV2023 guideline categories. **(G)** Confusion matrix depicting the performance of a classification model based on BD-EV quality control features.

We then investigated whether size distribution, EV protein abundance, and particle count could be used in machine learning to predict patient disease status. A OneVsRest classifier model was used to evaluate the abilities of the features described in Figure 1B-F to differentiate between disease classes. Since we have an imbalanced dataset for our classification model, we used the F1 score because it takes into account the different types of error-false positive and false negative, and not just number of incorrect predictions. The F1 score computes the average of precision and recall, where the relative contribution of both of these metrics results in the F1 score. In this first round of machine learning, the model could only reliably predict subjects based on the percentage of large EVs in the sample, or the number of proteins in the sample which pass a presence threshold in the case of PID (**Figure 1G**). This further corroborates the conclusion of the findings in Figure 1A-F that none of these features differ significantly between the patient groups except the ratio of small and large EVs in PID. This feature was not specific enough to differentiate between PiD and PSP, only CTRL from pathology (**Figure 1G**). Our findings show that tau pathology slightly affects BD-EV size and protein content, particularly in PID, but does not significantly impact their concentration or quality.

### 2 Mitochondrial / Endoplasmic Reticulum (ER) and Microtubule Protein Signatures in BD-EVs as Key Indicators of Tau Isoform Pathology

Given the large number of peptides present in BD-EVs, we first sought to determine whether protein profiles specific to 3R and 4R tauopathies were reflected in their content. To this end, we performed a Weighted Gene Co-Expression Network Analysis (WGCNA) to identify protein modules that best explained the variance between patient groups (**Figure 2A-B**). This approach revealed four significant modules: M1-Endoplasmic Reticulum, M2-Mitochondria, M3-Microtubules, and M4-Trivalent Inorganic Cation Transport (**Figure 2B**). Our results show that proteins associated with the endoplasmic reticulum and mitochondria are significantly more abundant in BD-EVs from patients with Pick’s disease (PiD) compared to controls (CTRL) and patients with PSP (Figure 2C). In contrast, microtubule-related proteins are significantly less abundant in PiD patients than in controls and PSP patients (**Figure 2C**). Using a predictive classification model suggests that only the M2-Mitochondria and M3-Microtubules modules enable good detection of PiD tauopathies, although they do not clearly distinguish between controls and PSP patients (**Figure 2D**).

**Figure 2.**
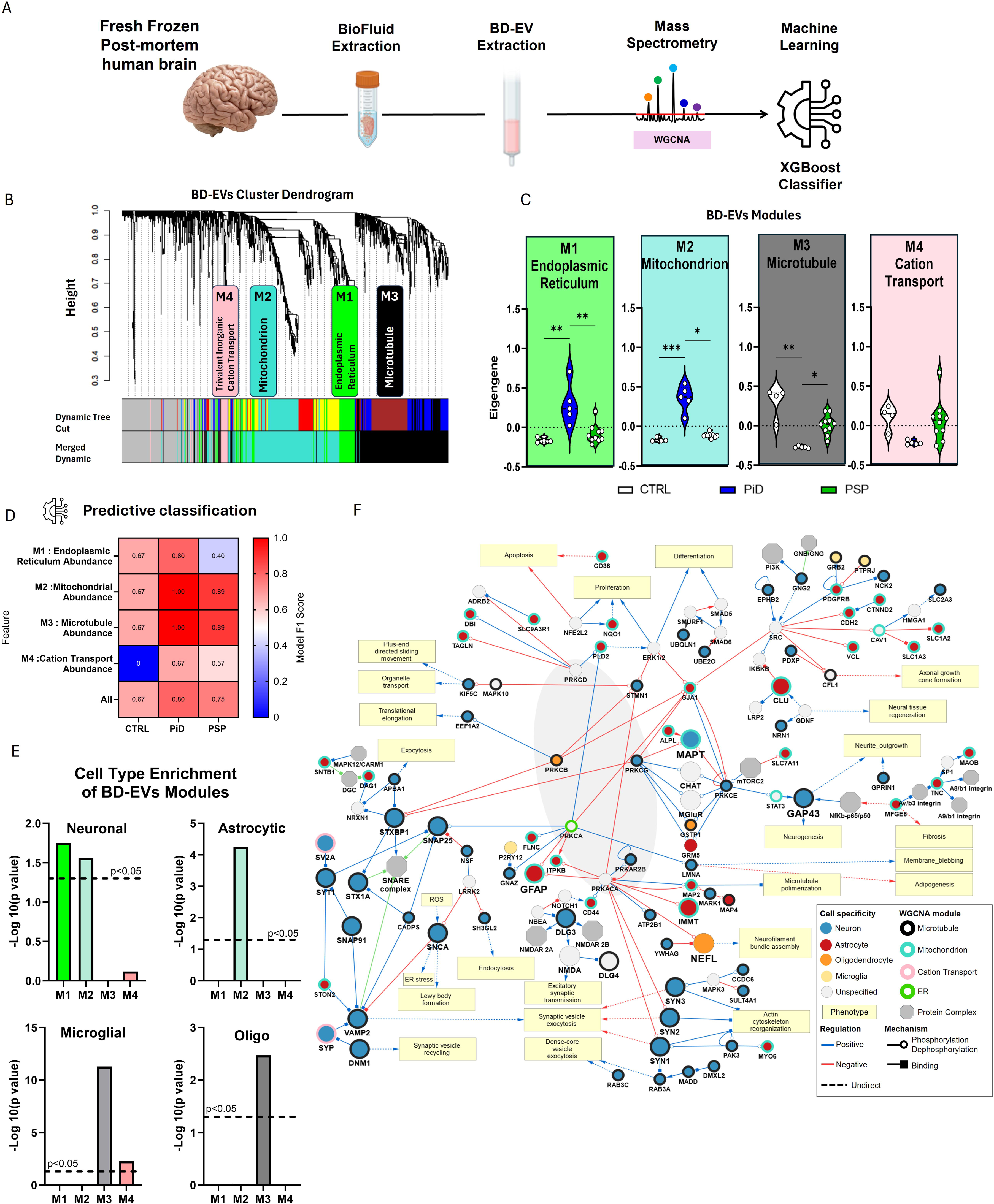
Distinct protein signatures of BD-EVs during 3R and 4R tauopathies. **(A)** Illustration of the procedure to investigate how BD-EVs may reflect brain tau pathology. Highlighted is the Weighted Gene Co-expression Network Analysis (WGCNA) of the BD-EV proteomic dataset. **(B)** Weighted gene co-expression network analysis emphasizing four protein modules which account for the majority of variance between patient groups: Endoplasmic Reticulum, Mitochondrion, Microtubule, and Trivalent Inorganic Cation Transport. **(C)** Violin plots of the cumulative abundance of the proteins contained within each of the four highlighted modules in WGCNA. **(D)** Confusion matrix depicting the performance of a classification model based on abundance of WGCNA module proteins. **(E)** Barplots representing brain cell type enrichment for corresponding WGCNA modules. We used -log10(p value) for FET values. Each barplot has a threshold with a single line for significant p value. **(F)** Protein-Protein Interaction Network of prospective biomarker proteins. Node colors are represents brain cell specificity: Blue: Neuron, Red: Astrocyte, Orange: Oligodendrocyte, Yellow: Microglia, Lightgrey: Unspecified; Node border color are representing the associated WGCNA module: Black: Microtubule module, Turquoise: Mitochondria Module, Pink: Cation Transport Module, Green: Endoplasmic Reticulum. Hexagonal node: Protein Complex, Rectangular Node: Phenotype. Edge Color represents type of regulation: Blue: Positive regulation, Red: Negative Regulation. Edge type represents mechanism: Open Circle: Phosphorylation and Dephosphorylation, Closed Square: Binding, Dashed line: Undirect interaction.

To further understand the origin and cell-type specificity of the proteins constituting BD-EVs, each protein was annotated using cell subtype-specific databases (see Methods). A Fisher’s exact test with Benjamini-Hochberg correction revealed that proteins in the Mitochondrion module are significantly associated with an astrocytic origin, while proteins in the Microtubule module are primarily of neuronal origin (**Figure 2E**). These results suggest that tau pathology may induce different perturbations depending on the cellular subtype involved. To deepen understanding of BD-EV content, we constructed a protein interaction network with SIGNOR. This interactome revealed that most proteins in the M2-Microtubule module (outlined in black) are neuron-derived (blue) and strongly interact with proteins from other modules (**Figure 2F**). Proteins in the M3-Microtubule module are known to play a key role in processes such as synaptic vesicle signalling, excitatory synaptic transmission, and synaptic vesicle recycling (e.g., SNAP25, SNAP91, STX1A, STX1B, SYT1, SV2A, DLG3, DLG4, VAMP2, DNM1, SNCA), as well as in neurofilament assembly, neurogenesis, actin cytoskeleton reorganization, axonal growth cone formation, and neurite outgrowth (GAP43, SYN1, SYN2, SYN3, PDXP). Conversely, most of the proteins in the M2-Mitochondrion module (outlined in turquoise) are astrocyte-derived (red) and interact closely with other proteins (IMMT, ITPKB, FLNC, GRM5, GFAP, CLU). These proteins are primarily involved in neuronal tissue regeneration and microtubule polymerization (**Figure 2F**).

In conclusion, our results demonstrate that the protein content of BD-EVs differs depending on whether the tauopathy is 3R or 4R. In Pick’s disease (3R), there is a higher abundance of astrocyte-derived mitochondrial proteins and a lower abundance of neuronal microtubule proteins compared to CTRL and PSP (4R) patients. These differences point to distinct cellular and functional alterations, for example in synaptic function, neuronal cytoskeletal structure, and astrocytic mitochondrial dynamics.

### 3 Tau Isoforms Influence the Neuronal and Glial Composition of BD-EVs

We investigated whether the type of tauopathy (3R or 4R) was reflected in the specific protein content of the BD-EV cell subtype (**Figure 3A**). We observed a significant decrease in the proportion of neuron-specific proteins in the PiD and PSP groups compared with the CTRL group (**Figure 3B**).

**Figure 3.**
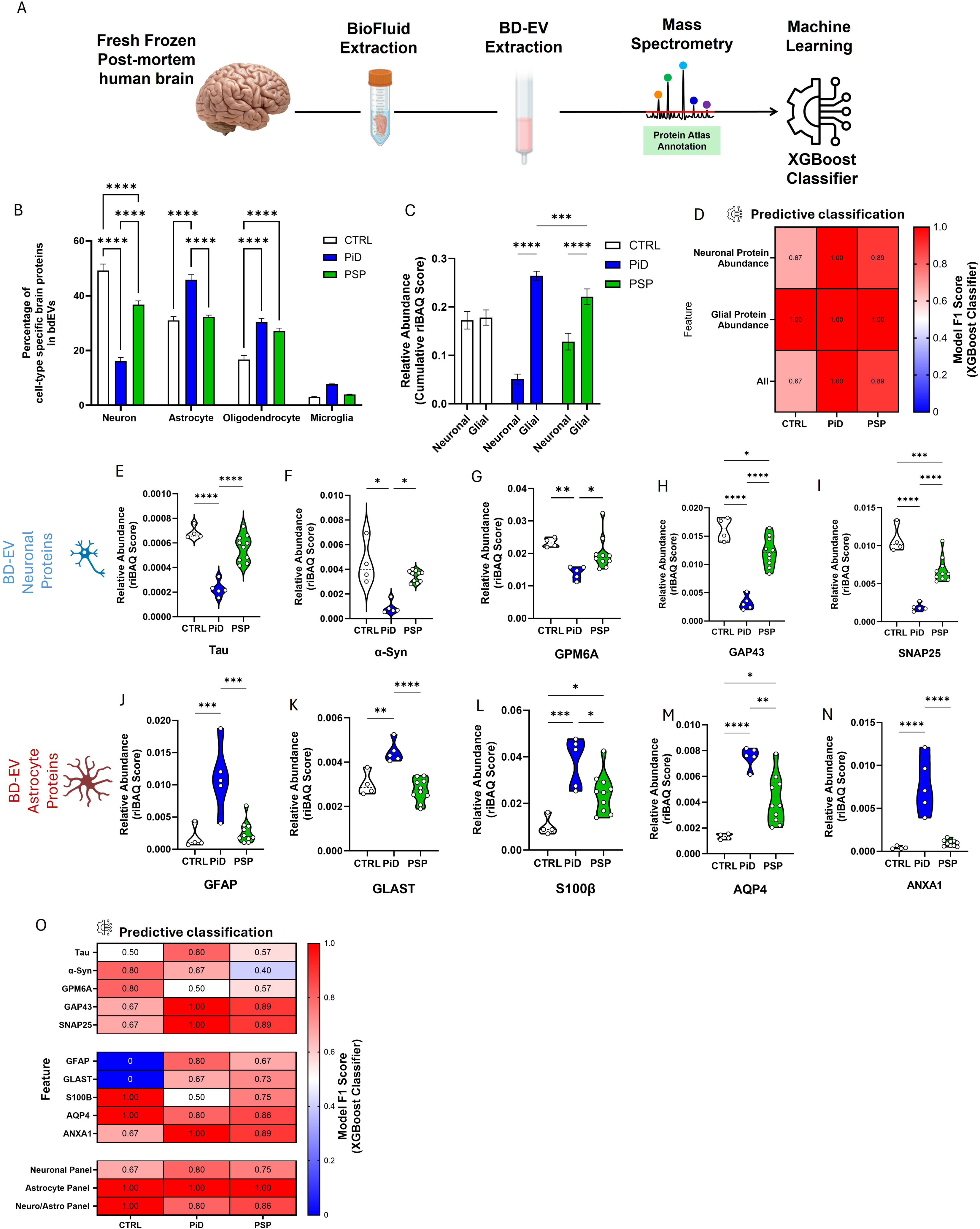
Neuronal and glial composition of BD-EV during 3R and 4R tauopathies. **(A)** Illustration of the procedure to investigate how BD-EVs may reflect brain tau pathology. Highlighted is the Human Protein Atlas annotation of the BD-EV proteomic dataset according to brain cell type specificity. **(B)** Bar graph showing the percentage of brain-specific proteins which can be categorized as neuronal, astrocytic, oligodendrocytic, or microglial per patient group. **(C)** Bar graph depicting the overall relative abundance of neuron-specific and glia-specific material by patient group. **(D)** Confusion matrix depicting the performance of a classification model based on abundance of glial and neuronal proteins. **(E-N)** Violin plots depicting the relative abundance of five neuronal (**E-I**) and astrocytic (**J-N**) proteins across patient groups. **(O)** Confusion matrix depicting the performance of a classification model based on abundance of specific neuronal and astrocytic proteins depicted in Fig. 3E-N.

At the glial level, the proportion of oligodendrocyte-specific proteins was significantly higher in the PiD and PSP groups, whereas the proportion of astrocyte-specific proteins was significantly increased only in 3R tauopathy (**Figure 3B**). Measurement of the relative amounts of neuronal and glial proteins (comprising astrocytes, microglia and oligodendrocytes) revealed a marked imbalance in 3R and 4R tauopathies (**Figure 3C**). Whereas the neuron/glial ratio is balanced at 1:1 in CTRL, this ratio has been radically altered in favour of glial enrichment in 3R and 4R patients, with a greater increase in glial proteins in 3R tauopathy (PID) compared with 4R (PSP), (**Figure 3C**). We then used the One-vs-Rest model to rank these different criteria and found that the sum of the relative amounts of glial proteins predicted CTRL, PID, and PSP with 100% accuracy (F1 score = 1), reflecting a perfect balance between precision and recall), (**Figure 3D**).

Among the many proteins associated with neuronal and glial categories, we selected five candidates that are abundant in BD-EV and relevant to the biomarker literature. For the neuronal proteins (**Figure 3E–I**), we observed that tau (**Figure 3E**), α-synuclein (**Figure 3F**), GPM6A (**Figure 3G**), GAP43 (**Figure 3H**), and SNAP25 (**Figure 3I**) were all significantly decreased in PiD. Although all of these proteins were significantly decreased in PiD compared to PSP, only GAP43 and SNAP25 showed a significant reduction when compared to the control group (**Figure 3E–I**).In contrast, astrocytic proteins (**Figure 3J-N**), such as GFAP (**Figure 3J**), GLAST, (**Figure 3K**), S100β (**Figure 3L**), AQP4 (**Figure 3M**), and ANXA1 (**Figure 3N**) were all significantly increased in PiD compared to controls and PSP patients. Using the machine learning model and the XGBoost classifier, we observed that among the selected neuronal proteins, certain proteins such as GAP43 and SNAP25 presented prediction scores > 0.89 to distinguish the 3R and 4R tauopathy groups. Astrocytic proteins such as AQP4 and S100b showed perfect prediction scores (F1 score = 1) to distinguish control patients from other pathologies (**Figure 3O**). Finally, our results show that combining the relative amounts of the five selected astrocytic proteins allows for a perfect differentiation (F1 score = 1) of 3R and 4R tauopathies from controls (CTRL), (**Figure 3O**). These results demonstrate that individual protein markers in BD-EV may be limiting for the purpose of distinguishing tauopathies, but that a combined analysis of glial proteins in BD-EV could significantly improve the diagnostic accuracy of 3R and 4R tauopathies.

### 4 BD-EV Protein Signatures: Correlation with Tau Pathology Severity

To determine whether BD-EV protein signatures from patients’ prefrontal cortex correlate with the severity of tau pathology in the same region, we analysed the relationships between the different BD-EV signatures and the quantification of AT8 staining previously performed on prefrontal cortex samples from the same patients, as reported by Leroux et al (**Figure 4A-B**). Among the neuronal proteins selected (**Figure 4C-G**), only a decrease in tau protein, GAP43 and SNAP25 showed a significant correlation with the increase in P-tau lesions (AT8) in this region. Conversely, the selected astrocytic proteins GFAP, GLAST, S100β, AQP4 and ANXA1 all showed a positive correlation with increased tau inclusions in patients’ brains (**Figure 4H-L**). However, although these correlations were significant, only GFAP demonstrated a high coefficient of determination (R² = 0.80), whereas the R² values for the other proteins remained relatively low (0.5 < R² >0.67). We then correlated the scores obtained previously (**Figures 1-3**) for the 3R and 4R signatures with the amount of AT8 staining observed in histological sections (**Figure 4A-B**). This analysis revealed that all the criteria studied showed a significant correlation, with high coefficients of determination (R²) for astrocytic protein abundance, neuronal protein abundance, and the mitochondrion module (**Figure 4M**). These results indicate that 3R tau pathology is associated with larger BD-EVs enriched in astrocytic material, mitochondrial residues, and endoplasmic reticulum material, but depleted in neuronal material (**Figure 4N**). This clear relationship between tau pathology severity and BD-EV characteristics suggests that these signatures could serve as valuable biological biomarkers for assessing tauopathy progression.

**Figure 4.**
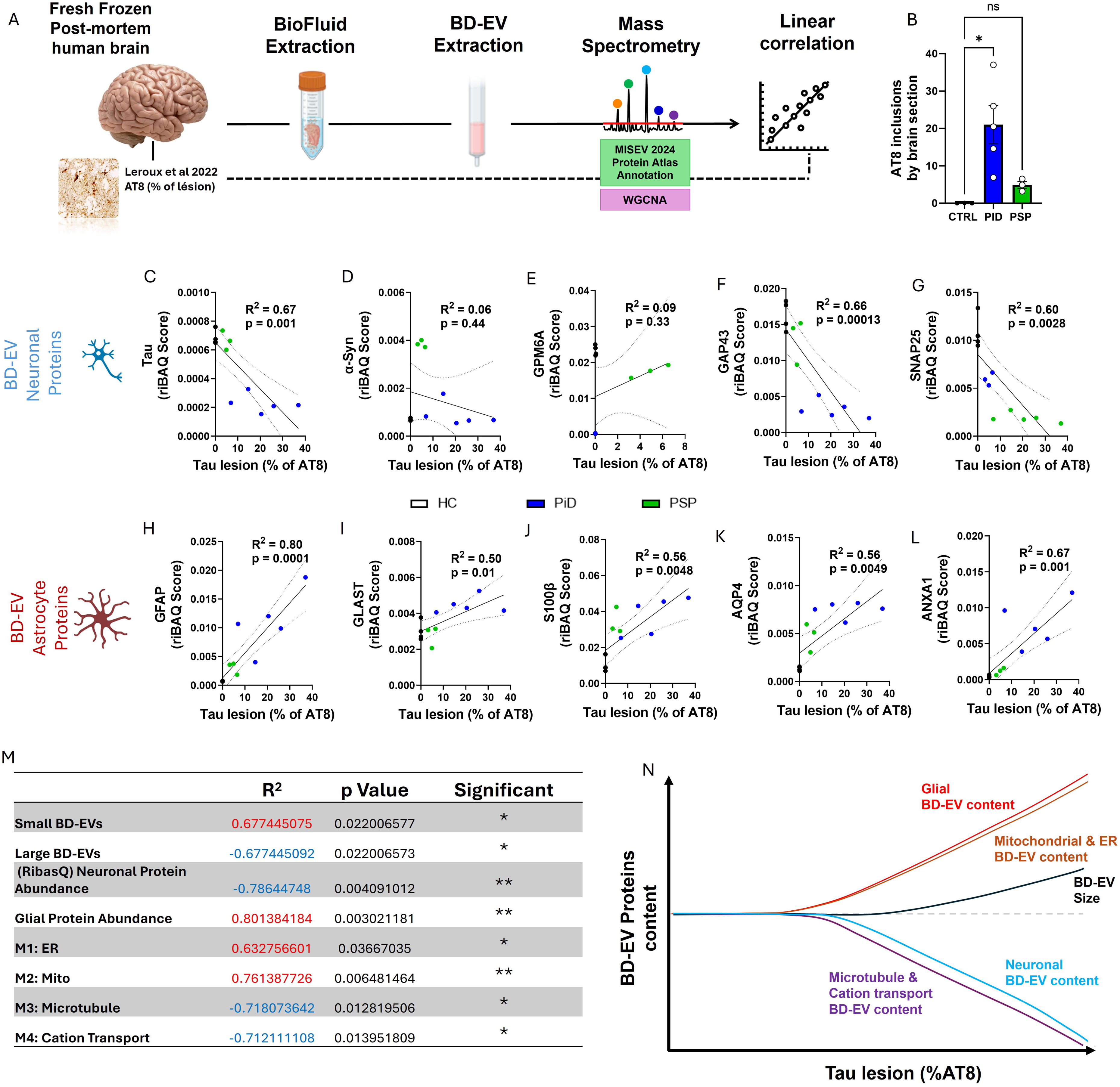
BD-EVs protein signatures are an indicator of Tau pathology severity. **(A)** Illustration of the procedure to investigate how BD-EVs may reflect brain tau pathology. Highlighted is the correlation of proteomic data with histopathological data from brain tissue of the same patients. **(B)** Bar graph showing the number of AT8 inclusions counted in brain slices by patient groups. **(C-L)** Scatter plots with simple linear regression depicting the correlation between the relative abundance of individual neuronal (**C-G**) and astrocytic (**H-L**) proteins with the percentage of AT8 tau lesions in the same patients. **(M)** Table showing the R2 values and p values for correlations between EV NTA and proteomic measurements with AT8 tau lesions in the same patients. **(N)** Hypothetical graph summarizing the evolution of BD-EV protein content as a function of Tau lesion severity (%AT8).

## Discussion

This study investigated the proteomic content of brain-derived extracellular vesicles (BD-EVs) isolated from the frontal cortex of patients with tauopathies and correlated these findings with the number and size of P-tau inclusions observed in this region through immunohistochemistry. Our results indicate that, although tau pathology does not significantly impact the concentration or quality of BD-EVs, it alters their size distribution and proteomic composition, particularly showing an enrichment in astrocytic and mitochondrial proteins in Pick’s disease. We also demonstrated that these distinct BD-EV signatures reflect the severity of tau pathology and could serve as potential biomarkers for monitoring tauopathy progression, highlighting the need for integrated proteomic approaches.

Although the field of biomarker discovery for neurodegenerative diseases, and particularly Alzheimer’s disease, has made considerable progress over the last decade, notably thanks to sophisticated tau proteomics methods and ultrasensitive P-tau 181 and P-tau 217 assays, it is still impossible to accurately differentiate between 3R and 4R tauopathies in the biological diagnostic setting ^14,15,42^. Indeed, PET tracers currently struggle to distinguish the two isoforms, and the use of circulating biomarkers runs up against a conceptual problem recently highlighted by Kyalu Ngoie Zola and colleagues ^17^. Indeed, although it is easy to classify 3R, 4R and mixed tauopathies in the insoluble fraction, this is not the case in the soluble fraction, suggesting that it will be difficult, despite the progress and precision of the tools, to imagine assaying 3R and 4R tau in biofluids ^17^. Our previous studies demonstrated that extracellular vesicles had very different consequences on the propagation of tauopathies and cell toxicity, particularly that of astrocytes ^43^. Here, we continue this work by analyzing the complete proteome of extracellular vesicles in patient brains, providing new insights and perspectives regarding extracellular vesicle diagnostics. We observed that pathology could influence the size of vesicles in the patient’s prefrontal cortex. However, our results show that the presence of tau aggregations, whether 3R or 4R, does not drastically change the concentration and quality of extracellular vesicles. We found a significant difference in the ratio of small (<150 nm) to large (>150 nm) vesicles in patients with PiD, which could indicate a change in the mechanism of vesicle secretion in non-apoptotic cells, or vesicles with a more extensive corona ^44^. Indeed, many free proteins could be secreted and adorn the corona of vesicles in the brain ^45^. In addition, large vesicles, derived from plasma membrane budding, may reflect alterations in cell biogenesis processes and cellular metabolism ^46^. Several studies have shown that pathological conditions, including certain brain diseases such as Alzheimer’s disease or cancer, can influence the size of the extracellular vesicles secreted ^47^.

To better understand the global protein signature of BD-EVs, we have examined the cellular signatures present in BD-EVs thanks to databases generated by McKenzie and colleagues ^32^. We found a significant difference in the neuronal to glial material ratio in pathological BD-EVs compared with control BD-EVs. Importantly, we also found a significant difference in the amount of glial material between PiD and PSP, indicating that the level of glial material in BD-EV may indicate the accumulation of specific tau isoforms in the brain. These results suggest that a central element between 3R and 4R tauopathies is probably how glial cells are differentially affected by the accumulation of the isoforms ^43,48,49^. Our previous work strongly suggested these results when we observed very different functional modifications by exposing astrocytes to EVs derived from neurons accumulating 3R or 4R tau ^43^. Many of our candidates detected as significant in BD-EVs correlate with the histopathological features of the same patients. Among them, the glial signature seems to be the most significant. This is one of the most critical aspects of this study, as it demonstrates that BD-EVs can reflect the molecular landscape of the brain affected by tau pathology. Interestingly, our other recent live-imaging study of the response of glial-neuronal 3D co-cultures exposed to extracellular paired helical filaments of tau (4R) revealed a strong astrocytic participation in the internalization of the aggregates ^50^, which may explain the glial signature in the results obtained here. Astrocytes (but not neurons) displayed increases in mitochondrial dynamics, reflecting the strong overlap between mitochondrial and astrocyte-specific proteins observed in the WGCNA analysis presented in this study. We also observed via a protein interaction network the regulatory role of astrocytic proteins from the mitochondrial module in different neuronal mechanisms such as synaptic transmission, neuronal cell differentiation and apoptosis, which highlights the importance of astro-neuro cross communication during tau pathology. Aside from numerous reported roles of astrocytes in the accumulation, processing, and response to tau ^43,51–53^, these observations reinforce the importance of this cell type, and most importantly its mitochondrial system, in the pathology and as target in research for highly specific biomarkers. Our study shows that the in-depth analysis of the BD-EV proteome offers new insights into the biological mechanisms underlying 3R and 4R tau aggregation. More importantly, we have shown that the amount of tau protein, alpha-synuclein, and glial signatures in BD-EVs reflect with excellent accuracy the histopathological state of the brain.

This study also demonstrated that the amount of tau protein peptides in BD-EVs is a good indicator of the type of tauopathy, 3R or 4R. Indeed, we observed a sharp decrease in tau in the BD-EVs of PiD relative to CTRL patients, making it possible to distinguish significantly between PiD and PSP patients, though not between PSP and CTRL patients. These results suggest that a simple assay of tau in BD-EVs could be indicative of 3R and 4R pathologies in the case where previous tests indicate the presence of a non-AD tau pathology. A more global analysis of molecular signatures by WGCNA shows that microtubule-associated proteins can differentiate PiD from both controls and PSP. This suggests that the difference between the effects of 3R and 4R accumulation on the cytoskeleton of cortical cells can be directly observed in BD-EVs. Several studies corroborate our hypothesis at the level of the protein in CSF. Indeed, p-tau181 and total tau in CSF are decreased in several tauopathies, particularly in comparison with AD. For example, FTLD-tau generally has lower levels of total tau and p-tau than AD ^14^, but patients with sporadic FTLD-TDP have even lower levels of p-tau181 than PSP, for which they could be used as diagnostic markers ^54,55^. Our results also show that the amount of alpha-synuclein protein follows the same trend as tau protein, i.e., it is inversely proportional to the levels of tau pathology in the brain. These results corroborate several studies that have demonstrated that neurons containing Pick’s bodies in the frontal cortex can also acquire Lewy bodies (LB) during disease progression ^56,57^. It has also been shown that in Alzheimer’s disease, neurons with tau aggregates are more likely to form LBs ^58^.

Furthermore, alpha-synuclein in neuron-derived EVs has recently been shown to be a potential biomarker of Parkinson’s disease (PD) ^59,60^. Compared with tau protein, alpha-synuclein is 10-fold more abundant in cortex BD-EVs. Our results therefore strongly suggest that measuring the number of tau peptides and alpha-synuclein in extracellular vesicles could be a good indicator for differentiating the progression and type of 3R and 4R tauopathies.

## Acknowledgements

The authors would like to thank Julien Gannevat from Nikon for his technical support, and Dr. Manfredo Quadroni from the proteomics platform at the University of Lausanne for his advice and technical support.

## Funding

The Swiss Confederation, Centre Hospitalier Universitaire Vaudois (CHUV), Synapsis Foundation, Lawson Foundation, and Marina Cuennet-Mauvernay Foundation financially supported this project. This work was also supported by grants from the program Investissement d’Avenir LabEx (investing in the future laboratory excellence), DISTALZ (Development of Innovative Strategies for a Transdisciplinary Approach to Alzheimer’s disease), ANR grants (TauSeed), and the PSP France Association. Our laboratories are also supported by LiCEND (Lille Centre of Excellence in Neurodegenerative Disorders), CNRS, Inserm, Métropole Européenne de Lille, the University of Lille, I-SITE ULNE, Région Hauts de France and FEDER.

## Competing Interests

The authors have no conflicts of interest to declare.

## Notes

### Competing Interest Statement

The authors have declared no competing interest.

